# Targeted chromosomal barcoding establishes direct genotype-phenotype associations for antibiotic resistance in *Mycobacterium abscessus*

**DOI:** 10.1101/2022.12.01.518803

**Authors:** Juan Calvet-Seral, Estefanía Crespo-Yuste, Vanessa Mathys, Hector Rodriguez-Villalobos, Pieter-Jan Ceyssens, Anandi Martin, Jesús Gonzalo-Asensio

## Abstract

A bedaquiline resistant *Mycobacterium abscessus* isolate was sequenced and a candidate mutation in the *atpE* gene was identified as responsible for the antibiotic resistance phenotype. To establish a direct genotype-phenotype relationship of this D29A mutation, we developed a recombineering-based method consisting of the specific replacement of the desired mutation in the bacterial chromosome. As surrogate bacteria, we used two *M. abscessus* antibiotic susceptible strains: ATCC19977, and the SL541 clinical isolate. The allelic exchange substrates used in recombineering carried either the sole D29A mutation, or a genetic barcode of silent mutations in codons flanking the D29A mutation. After selection of bedaquiline resistant *M. abscessus* colonies, transformed with both substrates, we obtained equivalent numbers of recombinants. These resistant colonies were analyzed by allele-specific PCR, and Sanger sequencing, demonstrating that the presence of the genetic barcode is linked to the targeted incorporation of the desired mutation in its chromosomal location. All recombinants displayed the same minimal inhibitory concentration to bedaquiline than the original isolate, from which the D29A mutation was identified. Finally, to demonstrate the broad applicability of this method, we confirmed the association of bedaquiline resistance with the *atpE* A64P mutation, performed in independent *M. abscessus* strains and by independent researchers.

## INTRODUCTION

The number of pulmonary infections caused by non-tuberculous mycobacteria (NTM) are increasing worldwide in the last decades [1], [2]. In 1987, the US center for disease control and prevention (CDC) estimated the NTM disease rate of 1.8/100.000 [3]. Data from North American studies between 2006 and 2012 suggested a disease rate of 5 to 10 per 100,000, which manifest the rising incidence in recent years [4]. Of those, the mayor cause of infections by fast growing mycobacteria is due to species from *Mycobacterium abscessus* complex, especially in patients with chronic lung diseases, such as cystic fibrosis or chronic obstructive disease [5], [6]. The *M. abscessus* complex is currently divided in three different subspecies *(abscessus, massiliense* and *bolletii)* in which *M. abscessus* subsp. *abscessus* (hereafter *M. abscessus)* is the most common pathogen [7]. Similarly to multidrug-resistant (MDR) tuberculosis, *M. abscessus* lung diseases are difficult to treat due to limited therapeutic options and its intrinsic and easy development of drug resistance [8], with an antimicrobial treatment success rate less than 50% [9], [10].

Different Health institutions recommend the use of a combination of oral macrolides (clarithromycin or azithromycin) with a parenteral β-lactams (cefoxitin or imipenem) and an aminoglycoside (amikacin) for a period of 6-12 months that can be extended to even 2 years [5]. These long-term treatments are usually poorly tolerated by patients, due to the manifestation of side effects and drug related toxicity against the different drugs used [10], [11]. In addition, *M. abscessus* can develop an inducible macrolide resistance conferred by the ribosomal methyltransferase gene *erm (41)* or acquire resistance due to mutations in 23S rRNA gene *rrl* [12]. This limited use of oral drugs makes nowadays essential the use of injectable antimicrobials. Therefore, there is a need to study new strategies to combat *M. abscessus* infections, especially for macrolide resistant strains. Several studies are evaluating the use of antimicrobials used in different mycobacterial infections like clofazimine [13], used to treat *M. leprae* and MDR tuberculosis; and bedaquiline [14] the first drug approved by the Food and Drug Administration in 40 years for treatment of MDR tuberculosis.

Bedaquiline is a diarylquinoline that inhibits the mycobacterial ATP synthase by targeting the subunit C of the F0/V0 complex, preventing rotor ring from acting as ion shuttle [15]. Sequencing of spontaneous resistant mutants obtained in the laboratory reveals that mutations in the coding gene of the ATPase subunit C, the *atpE* gene, are responsible of acquired resistant against bedaquiline in different mycobacterial species [16].

Emergence and spread of antimicrobial resistance is one of the top 10 global public health threats declared by the WHO [17]. As new mutations are daily identified in drug-resistant isolates, mainly thanks to lowering costs in the implementation of Whole Genome Sequencing technology, characterisation of these new mutations is essential to understand its impact in the bacterial biology, and specifically to corroborate their roles in antimicrobial resistance. This would allow to confirm the genetic-phenotype linkage of candidate mutations and discard other mechanisms like drug tolerance, based on phenotypic tolerability to drugs independent of a genetic association [18].

A classic strategy commonly used to confirm the genetic-phenotype association of a mutant resistant allele is the complementation and overexpression of the mutant allele in a sensitive strain with a plasmid. However, the resultant merodiploid strain carries both, mutant and sensitive alleles, which might lead to interfering phenotypes. In addition, overexpression of a gene product at non-physiological levels could also lead to exacerbated phenotypes which does not reflect a clinical situation. For example, the F0/V0 complex of bacterial ATPase is composed of several C subunits in which, if one of the subunits is susceptible of being blocked by bedaquiline, the whole rotor ring can be blocked. This case has been observed when trying to confirm two *atpE* mutations in *M. abscessus* (D29V and A64P), in which the expression of these mutant alleles of the *atpE* gene in a multicopy plasmid, did not increase the minimal inhibitory concentration (MIC) values against bedaquiline. However, when a selective pressure led to a spontaneous recombination replacing the whole sensitive allele in the *M. abscessus* chromosome, the resultant bacteria increased MIC 256 times [19].

Thus, a more optimal strategy to confirm genetic-phenotype associations in antimicrobial resistance, or whatever biological trait to be interrogated, implies to perform genetic manipulations directly in the bacterial chromosome. However, genetic tools for mycobacteria are limited, in comparison with widely used genetic model microorganisms as *Escherichia coli*, and genetic manipulation of *M. abscessus* is even more restricted. For example, some genome mutagenesis systems that work in other species of mycobacteria, like the thermosensitive-counterselective *sacB* system and the mycobacteriophage system, do not work in *M. abscessus* [20]. In the last years, suicide plasmids strategies based in counterselection with the *M. tuberculosis katG* sensitive allele or the *E. coli galK* gene have been successfully used to generate different *M. abscessus* knock-outs [21]–[24], even if construction of mutant strains is hardly correlated with antimicrobial resistance.

Recombineering system is one of the most used methods to perform genetic manipulations in the chromosome of mycobacteria. It was firstly described in *Escherichia coli* in 2000 [25] and later applied to *M. smegmatis* and *M. tuberculosis* [26]. It consists in the transformation of an allelic exchange substrate (AES) in a strain expressing the Gp60 and Gp61 proteins from the Che9c mycobacteriophage. Using a double strand AES (dsAES) with long homology flanking regions (around 1-1.5 Kbp) flanking an antibiotic resistant cassette, recombineering works with acceptable efficiency in *M. abscessus*. However, the percentage of double crossing-over mutants of *M. abscessus* is low in comparison with other mycobacteria due to the natural ability of *M. abscessus* to circularize lineal dsDNA fragments [20], [27], [28]. Recombineering system also allows the use of short simple strand oligonucleotides as AES (ssAES) in mycobacteria [29], however, as far as we known, there are no reports of the use of ssAES in *M. abscessus* to perform chromosome mutations.

In this work, we use the recombineering system to perform oligo-mediated targeted mutagenesis in the genome of two different strains of *M. abscessus*, the clinical strain SL541 and the laboratory reference strain *M. abscessus* ATCC19977, to confirm the *atpE* D29A sequenced mutation, which is involved in resistance to bedaquiline. In addition, we demonstrate the suitability of introducing barcoding, silent mutations to enable a fast and reliable colony screening by PCR-based methodologies, avoiding the extra time needed to confirm mutations by sequencing. This barcoding methodology has been successfully used to confirm *atpE* D29A mutation and, additionally, the *atpE* A64P, which also confers bedaquiline resistance in *M. abscessus*.

## MATERIALS AND METHODS

### Bacterial strains, culture media, and antibiotics

We worked with clinical isolate 81327881541, identified as *M. abscessus* by MALDI-TOF as described by Carbonnelle and colleagues [30] in the Cliniques universitaires Saint-Luc, Brussels, Belgium (*M. abscessus* SL541) and the laboratory reference strain *M. abscessus* ATCC19977 (GenBank accession number CU458896.1 [31]). Bacteria were cultured at 37°C in Middlebrook 7H9 medium (Difco) supplemented with 0.05% Tween-80 (Sigma) and 10% albumin-dextrose-catalase (ADC Middlebrook) in 25 cm^2^ flasks, without shaking. For plate growth, strains were incubated at 37°C in Middlebrook 7H10 (Difco) agar supplemented with 10% ADC Middlebrook. When required, Kanamicyn was added to *M. abscessus* at 50 μg/mL to maintain the pJV53 plasmid [26], and bedaquiline was added at a maximal concentration of 8 μg/mL.

### Isolation of a spontaneous mutant to bedaquiline

A spontaneous resistant mutant to bedaquiline was obtained from a single colony that grew after plating susceptible *M. abscessus* SL541 in 7H10-ADC containing 4 μg/mL of bedaquiline. This colony was subcultivated to stablish the *M. abscessus* SL541BQR strain, which was subjected to MIC determinations, DNA extraction and whole genome sequencing.

### Determination of MIC against *M. abscessus*

MICs in *M. abscessus* were determined on 7H10-ADC agar containing serial two-fold dilutions of bedaquiline ranging from 4 to 0.25 μg/mL, and in 7H9-ADC broth, bedaquiline was tested ranging from 8 to 0.0078 μg/mL. 10^5^ CFU were added to each well in agar plates and initial concentration of 10^5^ CFU/mL was used to determine MIC in broth. Plates were incubated at 37°C for 4 days. Then, 50 μL of 0.1 mg/mL filter-sterilized resazurin (Sigma-Aldrich) to agar plates, or 30 μL of resazurin or 2.5 mg/mL MTT [3-(4,5-dimethylthiazol-2-yl)-2,5-diphenyltetrazolium bromide] to broth plates. MIC was defined as the lowest concentration at which no color change was observed. Sensititre™ RAPMYCO Susceptibility Test was used as recommended by ThermoFisher to determine the antimicrobial resistance profile if *M. abscessus* SL541.

### Construction of recombinant mycobacteria and genetic manipulations

To prepare electrocompetent mycobacteria, 100 mL of bacterial culture was grown to an OD600 of 0.4-0.6. In the case of culturing bacteria with the recombineering system, acetamide was added to a final concentration of 0.2% (W/V) approximately one doubling time of the strain used before preparing the competent cells (3-4 hours) to allow the correct expression of the recombineering system. Bacterial pellets were washed several times in 10% glycerol-0.05% Tween-80 and finally resuspended in 1 mL of 10% glycerol. Aliquots of 100-200 μL were storage at −80°C for further use. The washing process was performed in cold conditions.

Aliquots of 100-200 μL electrocompetent cells were electroporated with 300-500 μg of plasmid DNA or 300-600 μg of the Allelic Exchange Substrate (AES) for recombineering (Table 1). 0.2 gap cuvettes (Bio-Rad) were used with a single pulse (2.5kV, 25μF, 1000Ω) in a GenePulser XcellTM (Bio-Rad). Cells were recovered with 5 mL of 7H9-ADC and incubated overnight at 37°C, to express the antibiotic resistance genes. Serial 10-fold dilutions were plated in 7H10-ADC plates containing the relevant antibiotic. Recombinant colonies typically grew in 8-10 days for *M. abscessus* and were tested by colony-PCR.

**Table 1.**
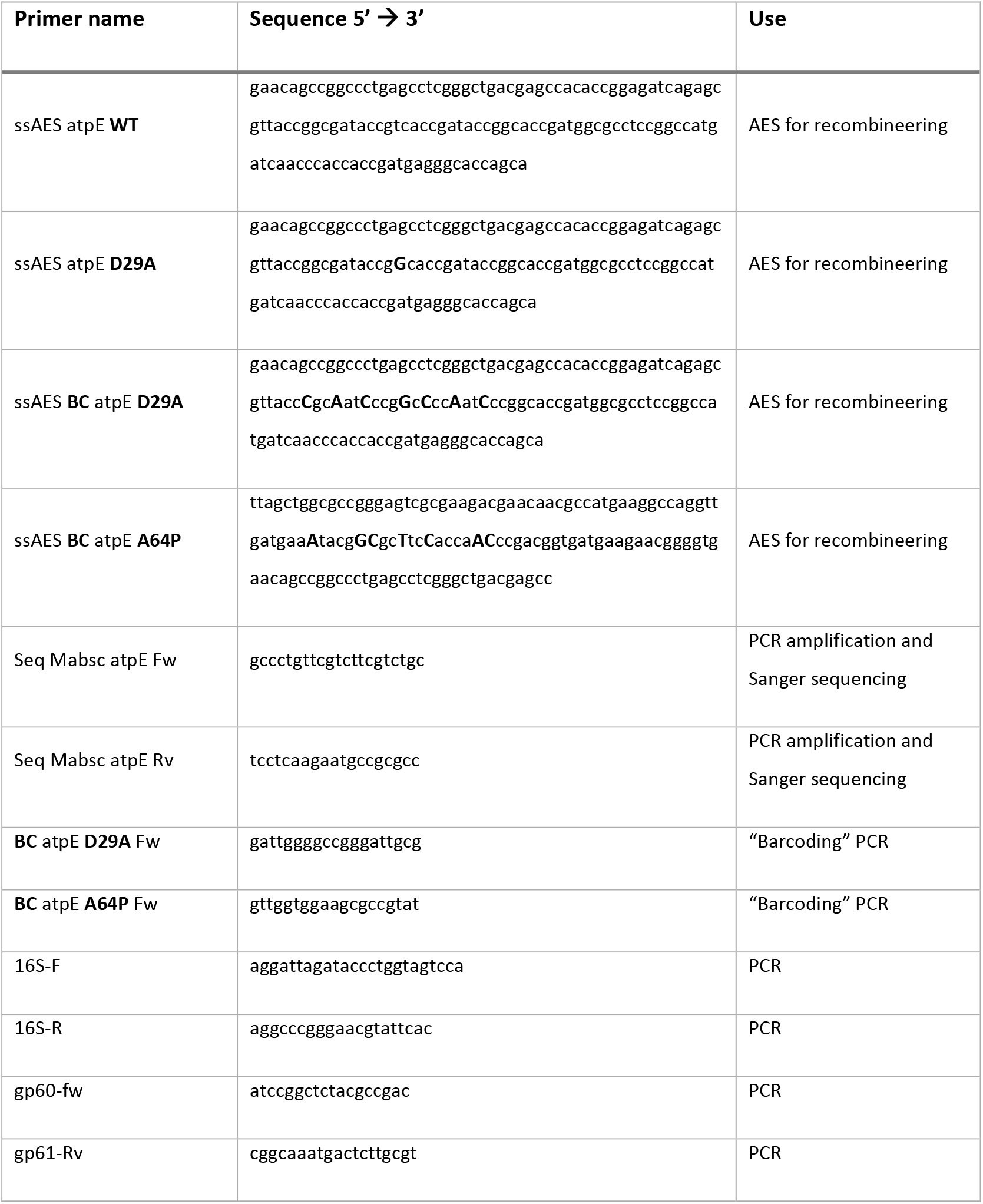
Oligonucleotides used for recombineering as AES and for PCR. Mismatching nucleotides introducing single point mutation (SNP) in the AES are highlighted in **bold**.

### Polymerase Chain Reaction and DNA sequencing

The genomic DNA from the different *M. abscessus* mutants was subjected to PCR amplification using primers from (Table 1). Bioline MyTaq DNA Polymerase was used for colony-PCR amplification with parameters as follows: heat denaturation at 95°C for 10 min followed by 30 cycles of 95°C for 15 seg, the corresponding annealing temperature for 15 seg and 72°C for 30 seg; and then a final extension at 72°C for 2 min. PCR products were visualized in agarose gels containing EtBr. PCR products for Sanger sequencing were treated with Affymetrix/Applied Biosystems ExoSAP-IT^®^ PCR Product Clean up according to instructions and sequenced by STAB VIDA Corp. to confirm mutations.

For Real-Time colony-PCR reaction, *M. abscessus* colonies were boiled colonies 15 min in 20 μL of sterilized water and centrifuged at maximal speed. TaKaRa TB Green^®^ Premix Ex Taq was used in a Step One Plus Real-Time PCR system with parameters as follows: heat denaturation at 95°C for 10 min followed by 40 cycles of 95°C for 10 seg, 62°C for 10 seg and 72°C for 50 seg. Primers for fast barcode-mutation detection were used at a final concentration of 0.25 μM and DNA of *M. abscessus* colonies was added from supernatant of boiled samples. Amplification of the specific Barcode-mutation was compared with amplification of 16s gene, as positive control of amplification.

For whole genome analysis, genomic DNA of *M. abscessus* strains under study was extracted and sequenced using Illumina MiSeq™ sequencing as described elsewhere (https://pubmed.ncbi.nlm.nih.gov/34751641/). Using CLC Genomics Workbench v20.0 (Qiagen), trimmed reads were either mapped against *M. abscessus* ATCC19977 or *de novo* assembled. Variant call in genes of interest was performed using CLC’s variant caller at high stringency, with minimal position coverage of 30x and minimal frequency 90%.

## RESULTS

### Identification of a bedaquiline resistant strain derived from a *M. abscessus* clinical isolate

The clinical isolate *M. abscessus* SL541 was obtained from a sputum sample from a patient with cystic fibrosis. The antimicrobial resistance profile of *M. abscessus* SL541 strains was determined with Sensititre™ RAPMYCO2 test, showing resistance to Cefoxitin, Imipenem, Tobramycin, Ciprofloxacin, Moxifloxacin, Trimethoprim, Sulfamethoxazole and Doxycycline (Table 2). Subsequently, a bedaquiline spontaneous resistant mutant was isolated from a single colony of *M. abscessus* SL541 plated in 7H10-ADC containing 4 μg/mL of bedaquiline. This clone, *M. abscessus* SL541BQR, showed a MIC agains bedaquiline in broth of 8-4 μg/mL, whereas its parental strain, *M. abscessus* SL541 exhibited a MIC of 0.25 μg/mL. The genomic DNA of both strains was isolated and sequenced using Illumina MiSeq, identifying the A89C single nucleotide mutation (SNP) in the *atpE* gene. This mutation translates into the non-synonymous D29A mutation in the AtpC subunit of the bedaquiline target ATP synthase, pointing as the causing mutation of bedaquiline resistance in our *M. abscessus* SL541 strain. Accordingly, we aimed to develop a genetic strategy to stablish a direct relationship between the genetic *atpE* mutation and the phenotypic resistance to bedaquiline in *M. abscessus* (Figure 1).

**Figure 1.**
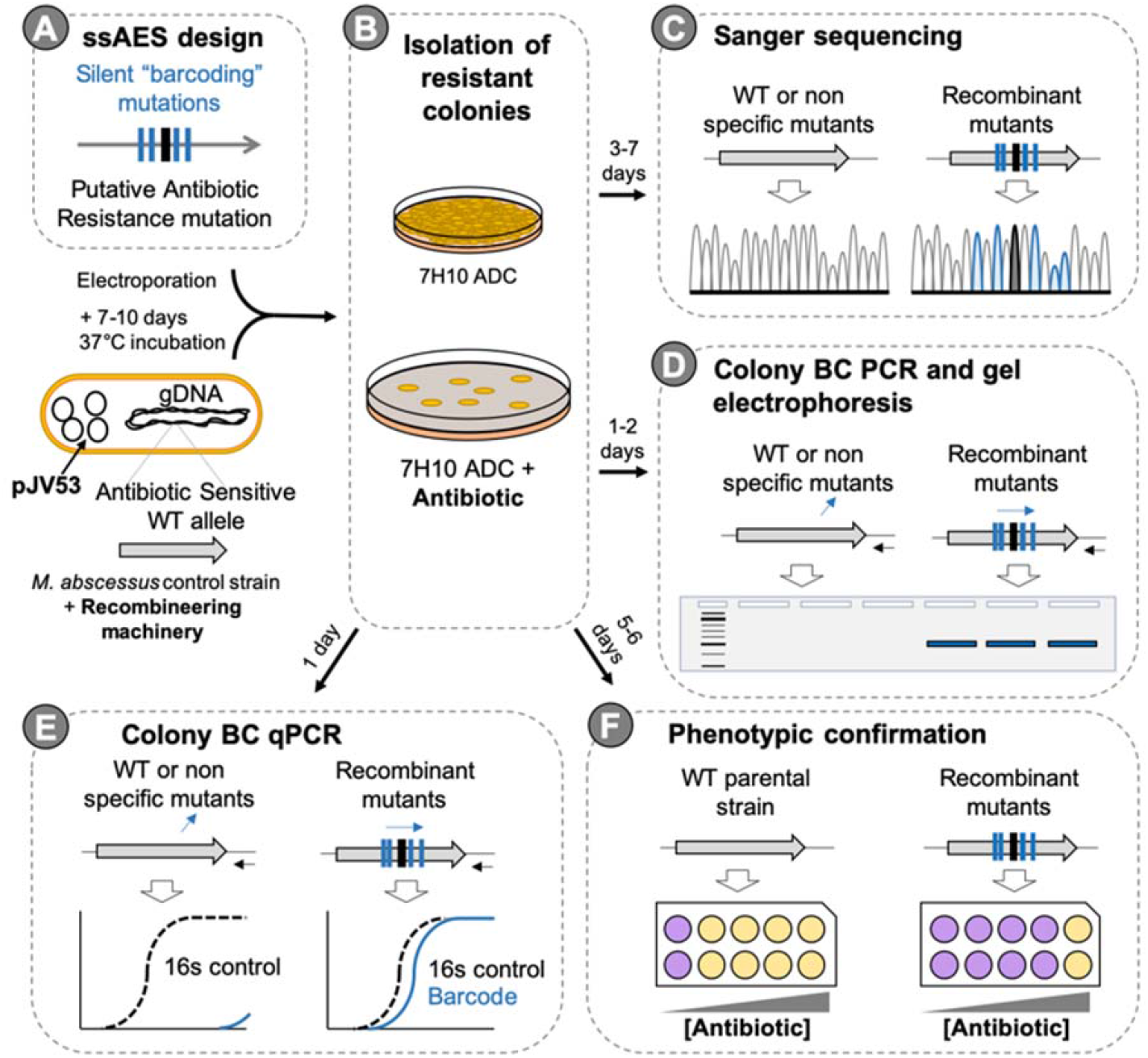
Workflow scheme followed in this study. (**A**) Design of ssAES carrying silent “barcoding” (BC) mutations and the antibiotic resistance mutation allow the (**B**) selection of resistant colonies after electroporation of *M. abscessus* carrying the recombineering machinery. (**C**) Sanger sequencing, (**D** and **E**) or Barcoding (BC)-PCR methods can be used to confirm genotype of recombinant colonies recovered. (**F**) Confirmation of the phenotype-genotype association of the mutation under study can be assessed by calculating the MIC to the antibiotic.

**Table 2.** Susceptibility profile of *M. abscessus* isolate 81327881541

| Antibiotics | *M. abscessus* clinical isolate 81327881541 |
| --- | --- |
| Amoxicillin/Clavulanate | > 64/32 ug/mL |
| Cefoxitin | R> 128 ug/mL |
| Ceftriaxone | > 64 µg/mL |
| Cefepim | > 32 µg/mL |
| Imipenem | R >64 µg/mL |
| Amikacin | = 16 µg/mL |
| Tobramycin | R > 16 µg/mL |
| Clarithromycin | = 0.12 µg/mL |
| Ciprofloxacin | R > 4 µg/mL |
| Moxifloxacin | R > 8 µg/mL |
| Trimethoprim/Sulfamethoxazole | R > 8/152 µg/mL |
| Minocycline | = 4 µg/mL |
| Doxycycline | R > 16 µg/mL |
| Tigecycline | = 2 µg/mL |

### Specific chromosomal replacements at the *atpE* 29^th^ codon position results in bedaquiline resistance in *M. abscessus*

Establishing a direct genotype-phenotype link in the context of antimicrobial resistance should follow molecular Koch’s postulates. Accordingly, the candidate gene (or mutation) responsible for drug resistance should be exclusively present in resistant, but not in susceptible bacteria. Then, introduction of the candidate gene (or mutation) into a susceptible bacterium would lead to development of drug resistance. However, introduction of a gene outside its original location might produce undesirable effects in the context of non-physiological expression of the mRNA or protein products. Further, introduction of genes using plasmids has two main disadvantages: on the one hand, this strategy usually leads to merodiploid bacteria which carry wild type and mutated copies of the gene of interest. On the other hand, plasmidic expression of genes does not resemble nor the copy number, nor the expression levels of chromosomal genes. With this scenario, we reasoned that an optimal strategy to confirm the role of a suspected gene (or mutation) in drug resistance, would require the replacement of the wild type allele in the chromosome. This strategy has the additional advantage that chromosomal replacements would eventually lead to drug resistant bacteria which could be selected in antibiotic containing plates.

We used single stranded recombineering, a technique which uses linear ssDNA substrates with flanking homology regions to direct targeted recombination in the chromosome [29]. We first confirmed the bedaquiline susceptibility of the *M. abscessus* SL541 strain in 7H10-ADC agar, in clear contrast to the *M. abscessus* SL541BQR carrying the bedaquiline resistance mutation *atpE* D29A (Figure 2A). We then introduced the recombineering machinery located in the pJV53 [26] plasmid into the *M. abscessus* SL541strain by electroporation. After selecting and confirming pJV53 positive clones, we confirmed that introduction of the recombineering plasmid did not alter the original bedaquiline susceptibility in the *M. abscessus* SL541 pJV53 strain (Figure 2A).

**Figure 2.**
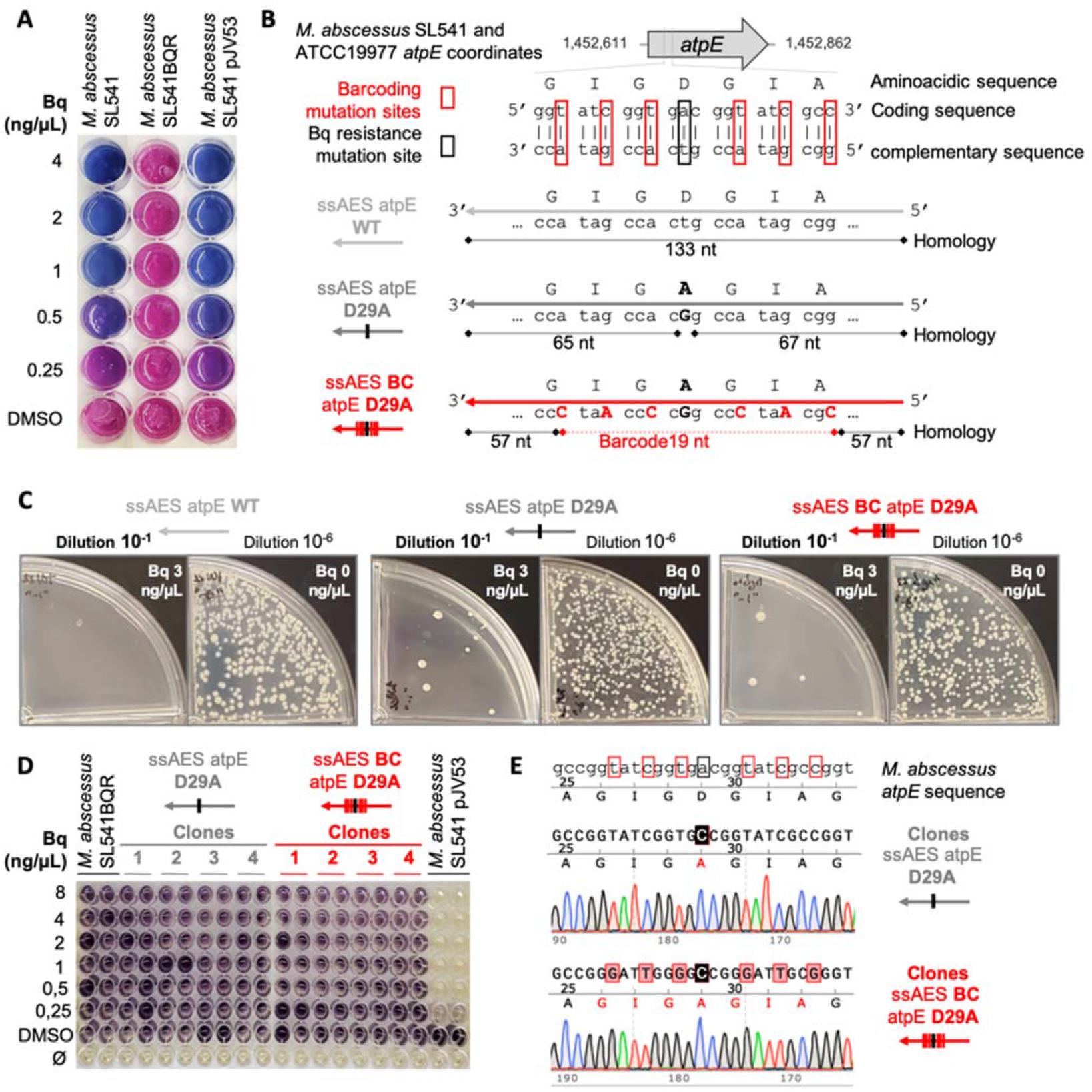
Chromosomal barcoding method to establish a genotype-phenotype association for the D29A mutation in the *atpE* gene. (**A**) Resazurin microtiter assay in 7H10-ADC to determine MIC of bedaquiline (Bq) against different *M. abscessus* strains. Wells with viable bacteria are shown in pink, whereas wells with absence of growth are shown in blue, after addition of resazurin. (**B**) Graphical representation of the ssAES used to mutate the 29^th^ codon of the *atpE* gene and their alignment in *M. abscessus* SL541 and reference ATTC19977 strains’ genome. (**C**) Electroporated bacteria with “ssAES atpE WT”, “ssAES atpE D29A” and “ssAES BC atpE D29A” were plated in presence (dilution 10^-1^) or absence (dilution 10^-6^) of Bq 3 μg/mL. (**D**) MTT in 7H9Tyl-ADC to determine the MIC of different isolated recombinant colonies. Wells with viable bacteria are shown in dark purple, whereas wells with absence of growth are shown in yellow, after addition of MTT. (**E**) Sanger sequencing chromatograms of recombinant colonies of *M. abscessus* electroporated with “ssAES atpE D29A” (up) and “ssAES BC atpE D29A” (down).

Once constructed the *M. abscessus* recombineering strain, we induced expression of the recombineering genes and then introduced allelic exchange substrates (AES) containing the desired mutation. Specifically, the AES consisted of single stranded oligonucleotides carrying either the wild type codon (GAC coding for Asp), or the mutated codon (GCC coding for Ala, and leading to a D29A mutation), in the central position of the ssDNA substrates. The left and right arms of the AES contained 65 and 67 homology nucleotides which promote the specific chromosomal replacement (Figure 2B). Recombinant bacteria were plated in medium containing bedaquiline, showing the presence of drug resistant colonies exclusively in bacteria transformed with the D29A allele, but not in transformants with the wild type allele (Figure 2C). To ensure that bedaquiline resistant bacteria arose from a similar number of transformants, we also plated bacteria without antibiotic. Results showed an equivalent number of CFUs in bacteria transformed with wild type and mutant alleles (Figure 2C). These bedaquiline resistant colonies were further confirmed by their increased MIC to bedaquiline relative to the *M. abscessus* recombineering strain (Figure 2D). Further, in order to discard that other mutations were contributing to our observed resistance phenotype, we sequenced the whole *atpE* gene in the recombinant colonies. We obtained a clear, and unique GCC substitution at the 29^th^ codon position relative to the wild type GAC codon in all analyzed colonies, indicative that this mutation is specifically responsible for the bedaquiline resistance phenotype in our recombinants (Figure 2E and Figure S1).

However, it should be noted that some of the *M. abscessus atpE* D29A mutants identified by this approach might arise by spontaneous mutations. Accordingly, we refined our strategy to specifically detect those resistant bacteria derived from our recombineering approach. We constructed a new AES containing the GCC mutated codon, immediately flanked by three silent mutations at each side, not altering the coding capacity of the mutated codons (Figure 2B). These silent modifications are aimed to act as a barcode to selectively detect the incorporation of the GCC mutation by recombineering, thus ruling out spontaneous mutations arisen during laboratory manipulation. This new AES contain 94.7% identical nucleotides relative to our previous approach (Figure 2B). Again, we obtained bedaquiline resistant colonies by applying this strategy (Figure 2C), which reproduced the antimicrobial resistance by elevated MIC to bedaquiline (Figure 2D). Interestingly, once sequenced the *atpE* gene, all bedaquiline resistant colonies contained the GCC alanine codon flaked by the barcode silent mutations (Figure 2E and S1). This barcoding strategy allowed to unequivocally demonstrate that introduction of the D29A mutation at the *atpE* chromosomal location by recombineering is responsible for bedaquiline resistance in the recombinants.

### Allele-specific PCR of bedaquiline resistant colonies allows the detection of chromosomal replacements at the *atpE* D29 codon position

Once demonstrated the utility of our genetic barcoding strategy to specifically detect the incorporation of the desired mutation by genomic sequencing, we prompted to optimize PCR-based methods to accelerate the whole process. First, we used allele-specific PCR, using an oligonucleotide annealing to the barcode sequence, but not to the wild type sequence, and a second oligonucleotide outside the ssDNA used as AES (Figure 3 A and B). After a PCR reaction, using a small portion of the recombinant colonies as genetic material, we confirmed the presence of the specific 531 bp PCR band in 10/12 bedaquiline resistant colonies transformed with the barcode oligonucleotide, but not in colonies transformed with the oligonucleotide containing solely the GCC mutation in the central position (Figure 3C). Overall, the whole process, from the transformation of the *M. abscessus* recombineering strain with the desired mutant AES, to the PCR verification of mutant colonies, requires less than two working weeks. Alternatively, we used Real Time-PCR to detect PCR amplification without the need for agarose gel electrophoresis. Results showed amplification of the 16S rRNA control gene (Ct≈25) in transformants with either the barcode, or the D29A only oligonucleotides (Figure 3D). However, using a D29A barcode-specific oligonucleotide, we were able to specifically detect amplification in our barcode transformants (Ct≈25) (Figure 3E), in contrast to the recombinant colonies transformed with the non-barcoded AES only containing the GCC mutation (Ct>31) (Figure 3D). We also confirmed the specificity of Real Time PCR products after interrogation of melting curves from each amplicon (Figures 3D and 3E). Using detection by Real Time PCR, the entire process lasted roughly 10 days.

**Figure 3.**
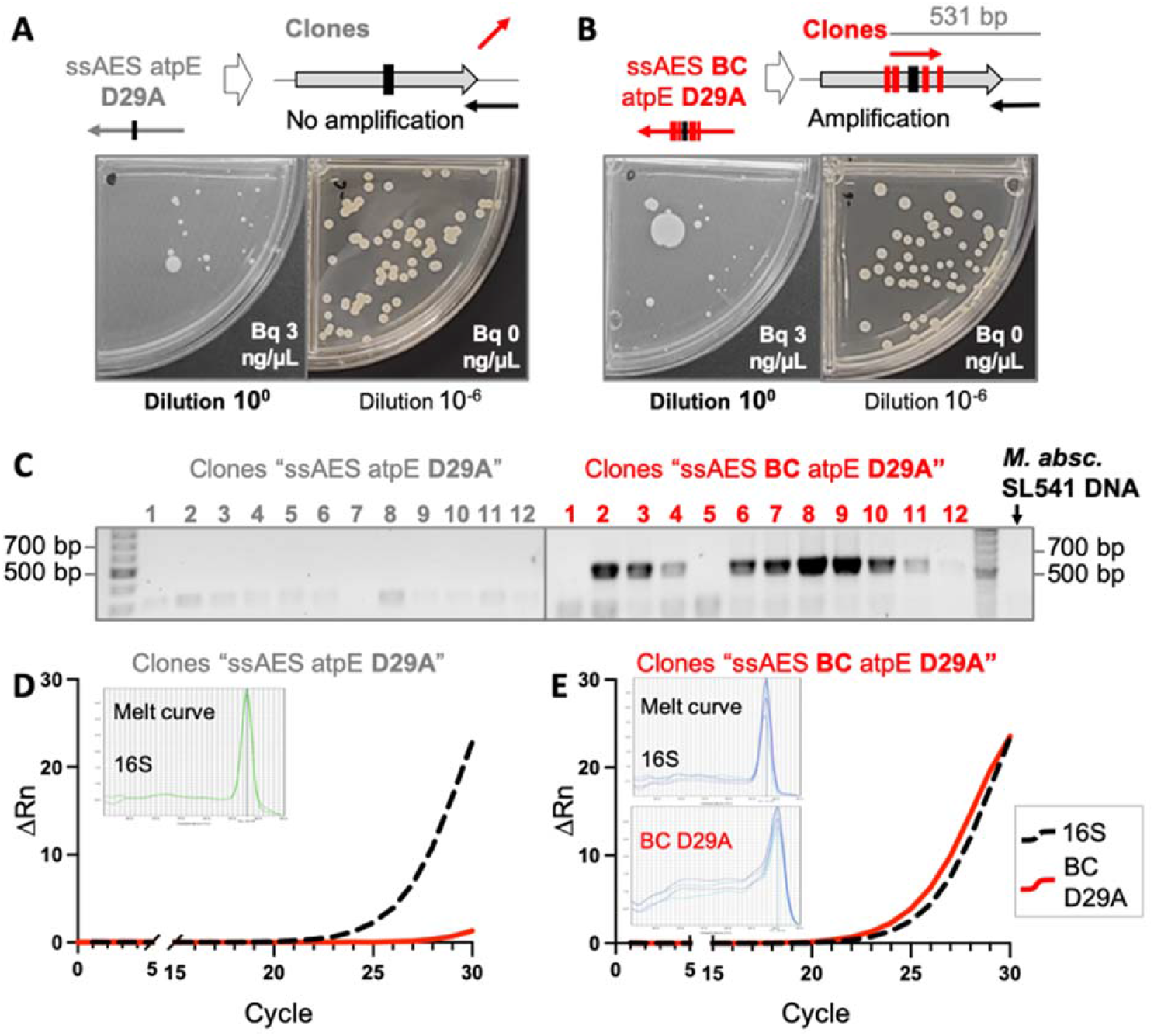
Development of PCR-based methods to identify chromosomal barcodes. (**A**) Graphical representation of the non-amplification and (**B**) amplification by barcode-PCR of bedaquiline (Bq) resistant bacteria recovered after electroporation with “ssAES atpE D29A” and “ssAES BC atpE D29A” respectively. (**C**) Agarose gels of barcoding-PCR of resistant colonies recovered. ΔRn values of amplification of the 16S RNA gene (dashed black) or the specific D29A barcode (continuous red) in (**D**) “ssAES atpE D29A” or (**E**) “ssAES BC atpE D29A”*M. abscessus* transformants obtained by barcode-RT-PCR.

In another attempt to demonstrate the robustness of our method, we aimed to reproduce the techniques described in this section with the *M. abscessus* ATCC19977 laboratory reference strain, carrying the pJV53 recombineering plasmid, in an experiment performed by an independent researcher. We successfully reproduced results obtained with either the barcode, or the non-barcode oligonucleotides (Figure S2A and B). Surprisingly, all *M. abscessus* bedaquiline resistant recombinants displayed a morphotype change from smooth to rough morphology, change not observed in recombinants from *M. abscessus* SL541 strain (Figure S2C). Altogether, our results demonstrate the robustness and reproducibility of our genetic replacement strategy, and lay foundations to apply these methods in independent laboratories.

### The barcoding genetic strategy was proven useful to detect D29A unrelated mutations in *M. abscessus*

Utilization of our genetic barcoding method as a general strategy to confirm antibiotic resistance mutations in *M. abscessus* requires confirmation with alternative polymorphisms. Accordingly, we selected an independent drug resistance mutation in *M. abscessus*. The A64P mutation has been described in *M. abscessus* [16], [19], and also in *M. tuberculosis*, [14], [16] as a polymorphism conferring bedaquiline resistance. We designed barcoded oligonucleotides carrying the 64^th^ Pro codon flanked by synonymous substitutions as described above (Figure 4A). Transformation of our *M. abscessus* SL541 recombineering strain with this AES, and culturing transformants in bedaquiline-containing plates, resulted in the growth of several drug resistant colonies (Figure 4B). Then, we used our PCR-based methods described above to confirm that specific incorporation of the barcode AES is responsible for the bedaquiline resistance of the transformants. By conventional, allele-specific PCR, we confirmed PCR amplification of the 432 bp specific band in all the assayed colonies (Figure 4C). We also confirmed specific Real Time PCR amplification of the barcoded *atpE* gene in recombinant (Ct≈18) (Figure 4E), but not in wild type (Ct>31) strains (Figure 4D). Again, the verification process lasted from 10 to 14 days, depending on the use of Real Time-PCR or conventional PCR, respectively. The bedaquiline resistance of the transformants was quantitatively confirmed by the MIC, which resulted >8 ng/⍰l, in contrast to the susceptible profile with their parental *M. abscessus* recombineering strain (Figure 4F).

**Figure 4.**
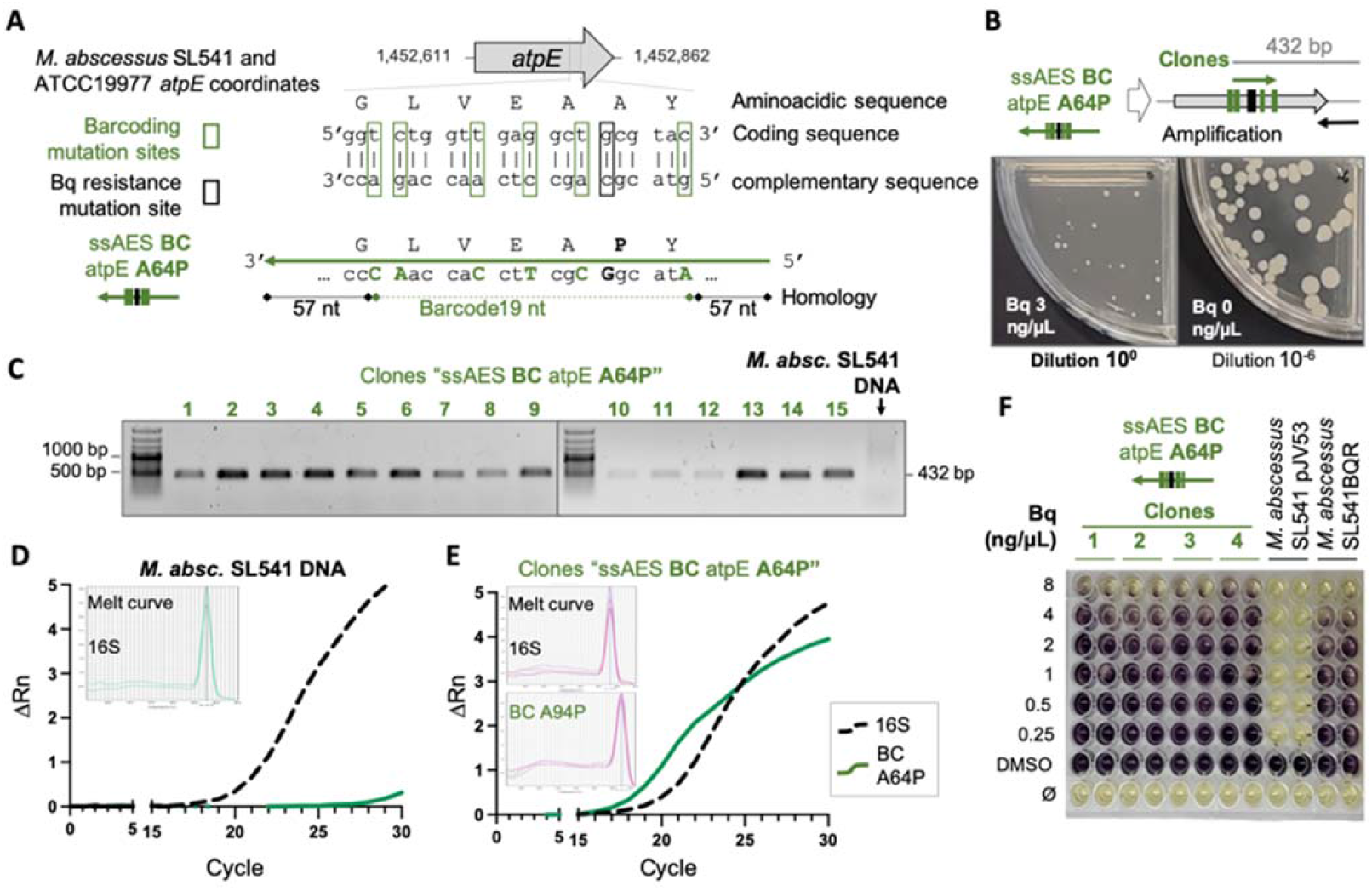
Chromosomal barcoding method to establish a genotype-phenotype association for the A64P mutation in the *atpE* gene. (**A**) Graphical representation of the ssAES used to mutate the 64^th^ codon of the *atpE* gene and their alignment in *M. abscessus* SL541 and reference ATTC19977 strains’ genome. (**B**) Graphical representation of the amplification by barcoding PCR of bedaquiline (Bq) resistant bacteria recovered after electroporation with “ssAES BC atpE A64P”. (**C**) Agarose gels of barcode-PCR of Bq resistant colonies. ΔRn values of amplification of the 16S RNA gene (dashed black) or the specific A64P barcode (continuous green) in (**D**) a “WT colony” or (**E**) “ssAES **BC** atpE **A64P**” *M. abscessus* transformants obtained by Barcode RT-PCR. (**F**) MTT in 7H9Tyl-ADC to determine the MIC of different isolated recombinant colonies. Wells with viable bacteria are shown in dark purple, whereas wells with absence of growth are shown in yellow, after addition of MTT.

## DISCUSSION

Antimicrobial resistance is a growing threat that endanger an effective treatment of infectious diseases and is indeed one of the health challenges for the 21^st^ century [17]. This is particularly important for mycobacterial infections since a limited repertoire of clinically approved drugs is available. Accordingly, mycobacterial infections usually require long term treatments with multiple drugs with the objective to reduce the chance of emergence of drug resistant strains. However, an inadequate adherence to treatment, prescription of inadequate drugs, or failure in drug supply, lead to antibiotic resistant mycobacteria. Drug resistance has been extensively studied in *M. tuberculosis* in the last years, which has led to an extensive catalogue of genome polymorphisms in the *inhA, katG*, and *rpoB* genes associated to resistance against isoniazid and rifampicin, the first line anti-tuberculous drugs [32]. Other antibiotic resistance polymorphisms have been also discovered for the first line drugs pyrazinamide and ethambutol, and, also, to second line drugs, which include bedaquiline and delamanid, recently incorporated to the anti-tuberculosis regimes [33], [34]. In the era of whole genome sequencing, these datasheets of antibiotic resistance polymorphisms are useful to delineate the possible drug resistance, or susceptibility profiles, by exploring genome data from clinical isolates. In this same context, PCR-based commercial systems to detect polymorphisms associated to drug resistance are widely available for *M. tuberculosis* [35]–[37]. However, these genomic methods have previously required establishing a direct link between chromosomal polymorphisms and antimicrobial resistance, and this has been otherwise the result of long-term observation and characterization of clinical isolates.

In this manuscript, we propose a genetic method aiming to accelerate the validation of drug resistance mutations in *M. abscessus*, a bacterium which similarly to drug-resistant *M. tuberculosis*, possess few therapeutic options [8]–[10]. We have focused here on bedaquiline resistance mutations, even though the use of this drug against *M. abscessus* infections was introduced in 2015 [38] and, accordingly, it is expected that bedaquiline resistance mutations are yet emerging after the recent introduction of the treatment. In addition, since the bedaquiline target is ATP synthase, whose *atp* coding genes show a high level of conservation between different mycobacteria (Figure S3), it is expected that results obtained with *M. abscessus* might be translated to related members of the *Mycobacterium* genus. Our method has been proven robust and reliable since it has been successfully tested with two unrelated mutations, in two independent *M. abscessus* strains, including a clinical isolate, and by independent researchers. This strategy presents several advantages with respect to other genetic methods: First, the specific chromosomal replacement avoids the use of ectopic gene expression, either in episomal or chromosomal locations. Second, the use of an available recombineering strain, and the synthesis of specific oligonucleotides, avoid the use of time-consuming cloning procedures. Third, results can be obtained in a reasonable short time, mostly limited to the growth of drug resistant colonies, which usually takes more than a week. Fourth, our strategy is scalable and allows simultaneous testing of various drug resistance genotypes. These advantages might be useful to deconvolve sequencing data related to drug resistant isolates. It is key to remember that interrogation of genome data does not always result in a list of polymorphisms located in well-known genes associated to drug resistance [32]. Accordingly, in these scenarios, each individual polymorphism should be genetically evaluated for its specific contribution to drug resistance. Additionally, as this barcoding strategy allows easily identification of our mutants, it can be optimized to compare different isogenic mutants and track how mutant populations evolve upon exposure to an antibiotic pressure, resembling signature-tagged mutagenesis, a technique used to distinguish desirable mutations in pools of transposon mutants in *M. tuberculosis* [39].

It remains to be answered whether the method described here is useful to detect low-level drug resistance, since we rely on the growth of drug resistant transformants on plates containing the corresponding drug. In this same context, further work is needed to ascertain whether other drug resistant mutations, aside from bedaquiline, can be tested by our method. However, regarding this latter observation, we should remember that our barcoding method is suitable to establish genotype-phenotype relationships in those polymorphisms located within coding regions, because of the need to introduce silent mutations in the vicinity of the targeted polymorphisms. Accordingly, it is not always possible to introduce such silent mutations in non-coding regions, as the ribosomal RNA subunits, which are the targets of aminoglycosides and macrolides used in the treatment of *M. abscessus* infections. These latter cases affecting non-coding RNA targets could be otherwise examined using non-barcoded oligonucleotides, and the subsequent verification of the gene sequence. Even though the non-barcoded method does not rule out the appearance of spontaneous mutations at the studied locus, a uniformity in the allele sequences from resistant colonies might be indicative of recombineering-derived drug resistant colonies. Indeed, in our hands, equivalent numbers of antibiotic resistant transformants were obtained when using non barcoded and barcoded AES (Figures 2 and 3), indicative that most, if not all, drug resistant colonies arose from AES recombination, and diminishing the chance of obtaining spontaneous drug resistant mutants.

Another aspect to consider is that silent mutations introduced in coding regions with this barcoding strategy could alter protein expression levels due to bacterial codon usage with the consequent change in bacterial fitness. It has been recently reported that *M. bovis* BCG is able to react to stress by tRNA reprograming and codon-biased translation. By this mechanism, BCG can drive the “over- or down-translation” mRNAs codon-biased from different families [40], [41]. Taking this into account, barcoding mutations could interfere in bacterial fitness if barcoded strains are used in different experiments (like *ex vivo* and *in vivo* infection experiments). mRNA structure and hybridization with other RNA structures of the bacteria could also be affected by barcoding mutations. However, this possible problem could be minimized with a rational design of the silent mutations. By maintaining percentage of codon usage, when feasible. On the other hand, *in silico* RNA analyzing servers are improving continuously and prediction of RNA structures and interactions could also be used to help in the design of our desired mutations.

Results described here could be theoretically applied to new candidate drugs which are currently in pre-licensing phases [42]. This would allow researchers to identify possible mechanisms of resistance prior to clinical evaluation of these forthcoming drugs, which might be useful to optimize diagnostic methods for the eventual drug resistant isolates. On the other side, since chromosomal replacements using recombineering have been described in different *Mycobacterium* species, including *M. tuberculosis, M. smegmatis, M. chelonae* [43]*, M canettii* (our unpublished results), we propose that our method could be transferable to other mycobacteria. This opens attractive possibilities to study not only drug resistance polymorphisms, but also metabolic, physiological, or virulence traits in *Mycobacterium*. Overall, our genetic strategy might help to accelerate the understanding the role of specific polymorphisms associated to drug resistance, with possible parallel application to understand mycobacterial biology.

## DATA AVAILABILITY

Data required to reproduce the content of this manuscript has been described elsewhere. In case of additional information, data will be provided upon request to the corresponding author.

## FUNDING

This work was supported by grants from “Gobierno de Aragón-Fondo Europeo de Desarrollo Regional (FEDER) 2014-2020: Construyendo Europa Desde Aragón” to J. C.-S.; Grant PRE2020-096507 funded by MCIN/AEI 10.13039/501100011033 to E.C.-Y.; and by Grant PID2019-104690RB-I00 funded by MCIN/AEI/ 10.13039/501100011033 to J. G.-A.

## ACKNOWLEDGEMENTS

Designed the study (J.-C.-S.; A.-M. and J. G.-A.), performed the experiments (J. C.-S.; E. C.-Y.; V. M. and P.-J. C.), analyzed the data (J. C.-S.; E. C.-Y.; V. M.; P.-J. C.; and J. G.-A.), provided biological material (H. R.-V. and A. M.), conceived figures (J.-C.-S. and J. G.-A.), designed figures (J.-C.-S.), wrote the original draft (J.-C.-S. and J. G.-A.), reviewed the final version of the manuscript (J.-C.-S. and J. G.-A.), obtained funding (J. G.-A.).

## DECLARATION OF COMPETING INTEREST

The authors declare that they have no known competing financial or personal interest relationships.

## ETHICS

The authors declare that, once published this manuscript, all drug resistant strains generated in this work will be properly eliminated.

## SUPPLEMENTARY FIGURES

**Figure S1.**
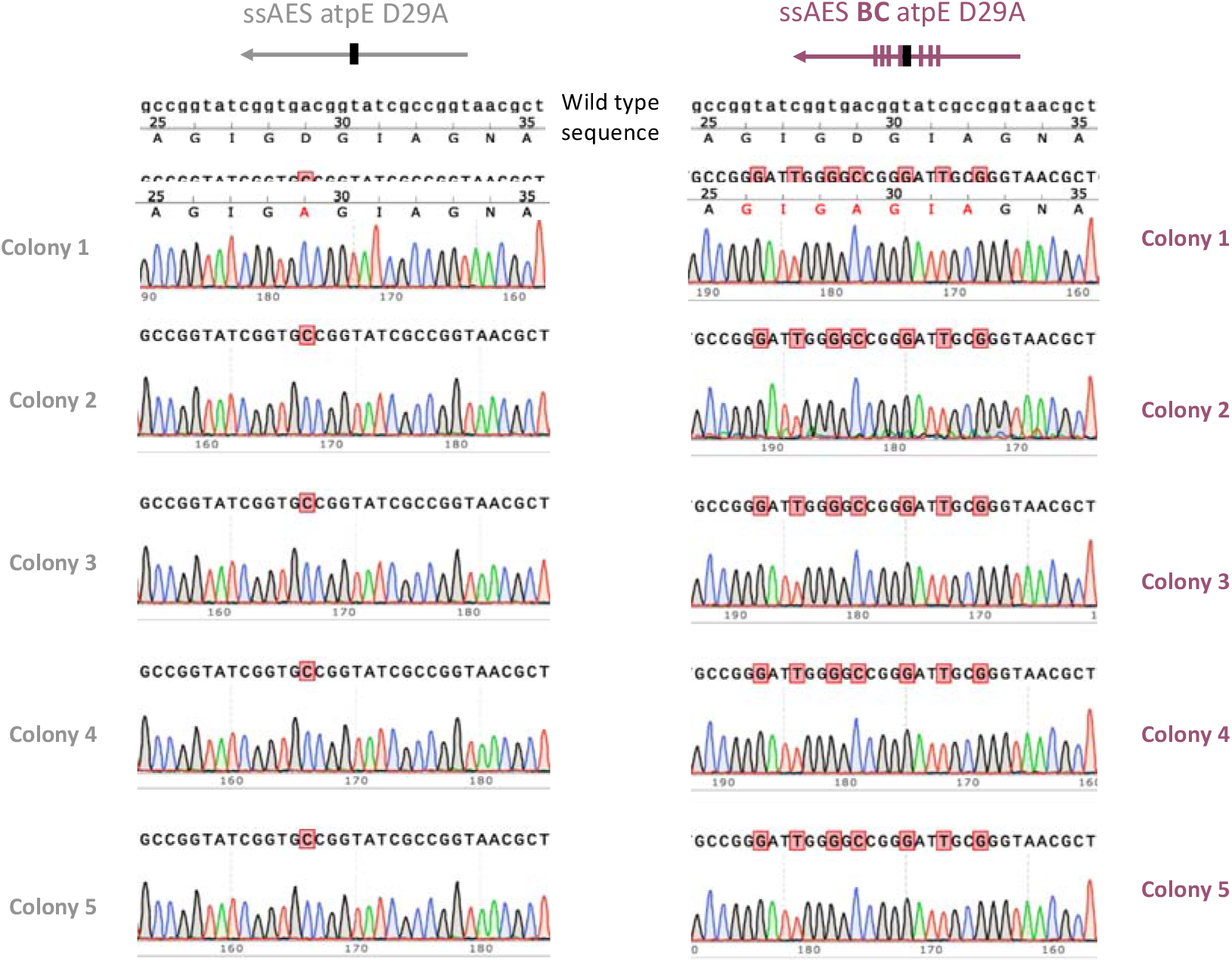
Sanger sequencing chromatograms of all recombinant colonies recovered of *M. abscessus* electroporated with “ssAES atpE **D29A**” (left) and “ssAES **BC** atpE **D29A**” (right).

**Figure S2.**
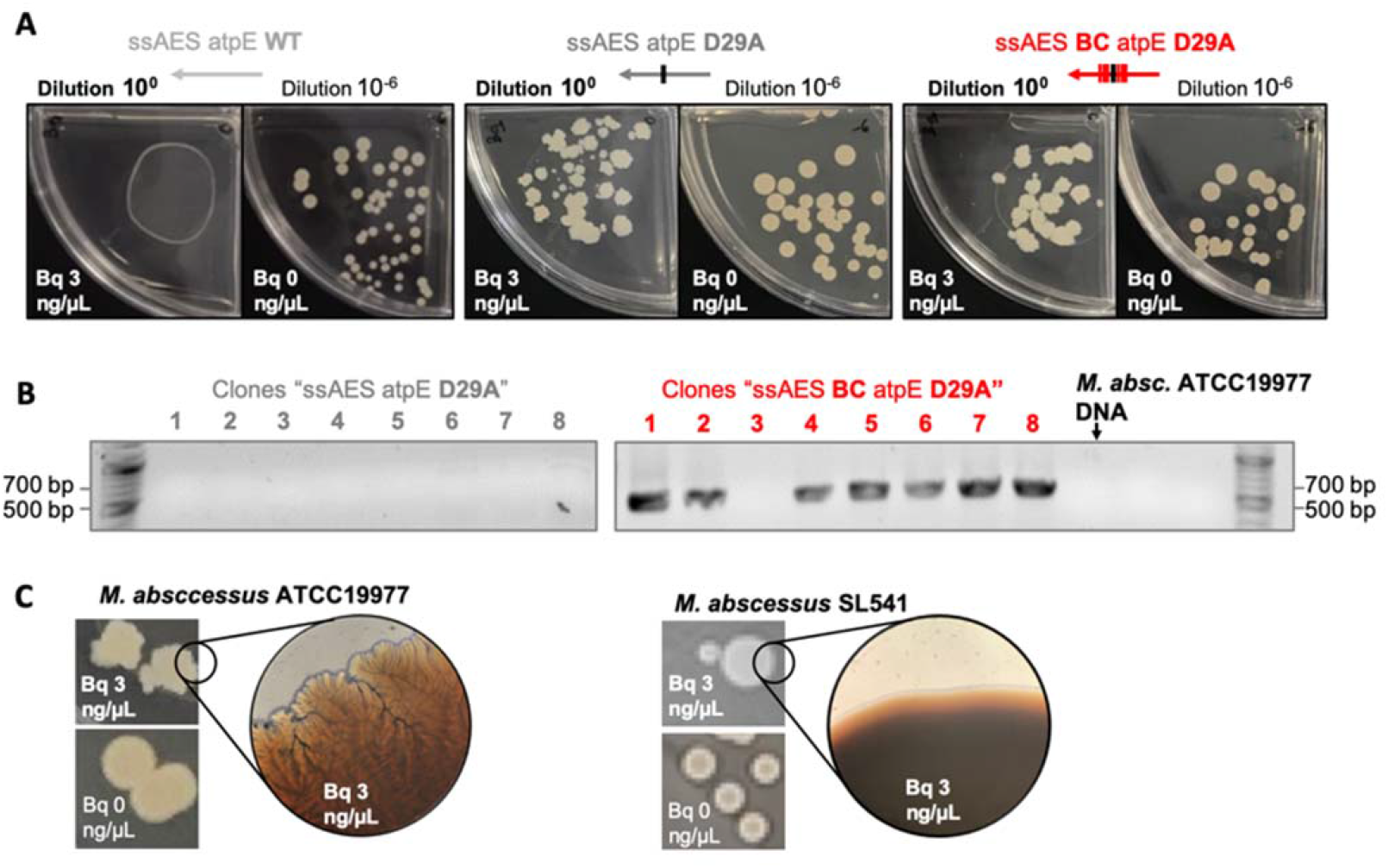
(**A**) Electroporated *M. abscessus* pJV53 (UZ) with “ssAES *atpE* WT”, “ssAES *atpE* D29A” and “ssAES BC *atpE* D29A” was plated in absence (dilution 10^-6^) or presence (dilution 10^0^) of bedaquiline (Bq) at 3 μg/mL. (**B**) Resistant colonies were subjected to Barcode-PCR. (**C**) Morphology of recombinant colonies obtained in presence of Bq 3 μg/mL from *M. abascessus* ATCC19977 changed from smooth to rough. No change is observed in *M. abscessus* SL541 recombinants.

**Figure S3.**
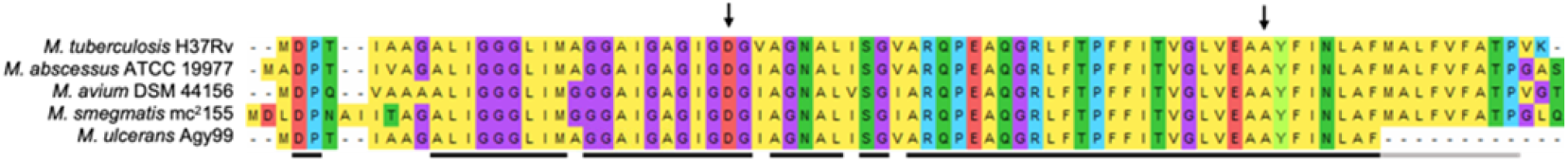
Alignment by MUSCLE algorithm in MEGA software of *atpE* encoded protein (ATP synthase subunit C) of different mycobacteria. Residues mutated in this study are marked with arrows. 100% identity regions are underlined, showing high conservation of the protein (black line, conserved in the five species compared, grey line, conserved in all but deleted in *M. ulcerans* Agy99 strain).

## REFERENCES

[1] C. N. Ratnatunga et al., ‘The Rise of Non-Tuberculosis Mycobacterial Lung Disease’, Frontiers in Immunology, vol. 11. Frontiers Media S.A., p. 303, Mar. 03, 2020. doi: 10.3389/fimmu.2020.00303.

[2] J. E. Stout, W. J. Koh, and W. W. Yew, ‘Update on pulmonary disease due to non-tuberculous mycobacteria’, International Journal of Infectious Diseases, vol. 45. Elsevier, pp. 123–134, Apr. 01, 2016. doi: 10.1016/j.ijid.2016.03.006.

[3] B. A. Kendall and K. L. Winthrop, ‘Update on the epidemiology of pulmonary nontuberculous mycobacterial infections’, Semin Respir Crit Care Med, vol. 34, no. 1, pp. 87–94, 2013, doi: 10.1055/S-0033-1333567.

[4] D. R. Prevots and T. K. Marras, ‘Epidemiology of human pulmonary infection with nontuberculous mycobacteria a review’, Clinics in Chest Medicine, vol. 36, no. 1. NIH Public Access, pp. 13–34, Mar. 01, 2015. doi: 10.1016/j.ccm.2014.10.002.

[5] R. A. Floto et al., ‘US Cystic Fibrosis Foundation and European Cystic Fibrosis Society consensus recommendations for the management of non-tuberculous mycobacteria in individuals with cystic fibrosis’, Thorax, vol. 71, no. Suppl 1, pp. i1–i22, Jan. 2016, doi: 10.1136/THORAXJNL-2015-207360.

[6] D. E. Griffith et al., ‘An Official ATS/IDSA Statement: Diagnosis, Treatment, and Prevention of Nontuberculous Mycobacterial Diseases’, https://doi.org/10.1164/rccm.200604-571ST, vol. 175, no. 4, pp. 367–416, Dec. 2012, doi: 10.1164/RCCM.200604-571ST.

[7] M. D. Johansen, J. L. Herrmann, and L. Kremer, ‘Non-tuberculous mycobacteria and the rise of Mycobacterium abscessus’, Nature Reviews Microbiology, vol. 18, no. 7. Nature Publishing Group, pp. 392–407, Feb. 21, 2020. doi: 10.1038/s41579-020-0331-1.

[8] R. Nessar, E. Cambau, J. M. Reyrat, A. Murray, and B. Gicquel, ‘Mycobacterium abscessus: A new antibiotic nightmare’, Journal of Antimicrobial Chemotherapy, vol. 67, no. 4, pp. 810–818, 2012, doi: 10.1093/jac/dkr578.

[9] N. T. Quang and J. Jang, ‘Current Molecular Therapeutic Agents and Drug Candidates for Mycobacterium abscessus’, Front Pharmacol, vol. 12, no. August, 2021, doi: 10.3389/fphar.2021.724725.

[10] L. Victoria, A. Gupta, J. L. Gómez, and J. Robledo, ‘Mycobacterium abscessus complex: A Review of Recent Developments in an Emerging Pathogen’, Frontiers in Cellular and Infection Microbiology, vol. 11. Frontiers Media S.A., p. 338, Apr. 26, 2021. doi: 10.3389/fcimb.2021.659997.

[11] J. Lyu et al., ‘Outcomes in patients with Mycobacterium abscessus pulmonary disease treated with long-term injectable drugs’, Respir Med, vol. 105, no. 5, pp. 781–787, May 2011, doi: 10.1016/j.rmed.2010.12.012.

[12] S. J. Shallom, N. S. Moura, K. N. Olivier, E. P. Sampaio, S. M. Holland, and A. M. Zelazny, ‘New real-time PCR assays for detection of inducible and acquired clarithromycin resistance in the mycobacterium abscessus group’, J Clin Microbiol, vol. 53, no. 11, pp. 3430–3437, Nov. 2015, doi: 10.1128/JCM.01714-15.

[13] H. Choi et al., ‘Clinical characteristics and treatment outcomes of patients with acquired macrolide-resistant Mycobacterium abscessus lung disease’, Antimicrob Agents Chemother, vol. 61, no. 10, 2017, doi: 10.1128/AAC.01146-17.

[14] K. Andries et al., ‘A Diarylquinoline Drug Active on the ATP Synthase of Mycobacterium tuberculosis’, Science (1979), vol. 307, no. 5707, pp. 223–227, Jan. 2005, doi: 10.1126/science.1106753.

[15] L. Preiss et al., ‘Structure of the mycobacterial ATP synthase Fo rotor ring in complex with the anti-TB drug bedaquiline’, Sci Adv, vol. 1, no. 4, 2015, doi: 10.1126/sciadv.1500106.

[16] E. Segala, W. Sougakoff, A. Nevejans-Chauffour, V. Jarlier, and S. Petrella, ‘New mutations in the mycobacterial ATP synthase: New insights into the binding of the diarylquinoline TMC207 to the ATP synthase C-Ring structure’, Antimicrob Agents Chemother, vol. 56, no. 5, pp. 2326–2334, 2012, doi: 10.1128/AAC.06154-11.

[17] ‘Antimicrobial resistance’. https://www.who.int/news-room/fact-sheets/detail/antimicrobial-resistance (accessed Oct. 17, 2022).

[18] E. M. Windels, B. van den Bergh, and J. Michiels, ‘Bacteria under antibiotic attack: Different strategies for evolutionary adaptation’, PLoS Pathog, vol. 16, no. 5, p. e1008431, May 2020, doi: 10.1371/journal.ppat.1008431.

[19] C. Dupont et al., ‘Bedaquiline inhibits the ATP synthase in mycobacterium abscessus and is effective in infected zebrafish’, Antimicrob Agents Chemother, vol. 61, no. 11, Nov. 2017, doi: 10.1128/AAC.01225-17.

[20] H. Medjahed and J. M. Reyrat, ‘Construction of mycobacterium abscessus defined glycopeptidolipid mutants: Comparison of genetic tools’, Appl Environ Microbiol, vol. 75, no. 5, pp. 1331–1338, 2009, doi: 10.1128/AEM.01914-08.

[21] A. Rominski, P. Selchow, K. Becker, J. K. Brülle, M. Dal Molin, and P. Sander, ‘Elucidation of Mycobacterium abscessus aminoglycoside and capreomycin resistance by targeted deletion of three putative resistance genes’, Journal of Antimicrobial Chemotherapy,vol. 72, no. 8, pp. 2191–2200, Aug. 2017, doi: 10.1093/jac/dkx125.

[22] S. A. Gregoire, J. Byam, and M. S. Pavelka, ‘GalK-based suicide vector mediated allelic exchange in mycobacterium abscessus’, Microbiology (United Kingdom), vol. 163, no. 10, pp. 1399–1408, Oct. 2017, doi: 10.1099/mic.0.000528.

[23] A. Viljoen, A. V. Gutiérrez, C. Dupont, E. Ghigo, and L. Kremer, ‘A simple and rapid gene disruption strategy in Mycobacterium abscessus: On the design and application of glycopeptidolipid mutants’, Front Cell Infect Microbiol, vol. 8, no. MAR, Mar. 2018, doi: 10.3389/fcimb.2018.00069.

[24] J. M. Belardinelli et al., ‘Therapeutic efficacy of antimalarial drugs targeting DosRS signaling in Mycobacterium abscessus’, Sci Transl Med, vol. 14, no. 633, p. 3860, 2022, [Online]. Available: https://www.science.org/doi/abs/10.1126/scitranslmed.abj3860

[25] K. A. Datsenko and B. L. Wanner, ‘One-step inactivation of chromosomal genes in Escherichia coli K-12 using PCR products’, Proc Natl Acad Sci U S A, vol. 97, no. 12, pp. 6640–6645, 2000, doi: 10.1073/pnas.120163297.

[26] J. C. van Kessel and G. F. Hatfull, ‘Recombineering in Mycobacterium tuberculosis’, Nat Methods, vol. 4, no. 2, pp. 147–152, 2007, doi: 10.1038/nmeth996.

[27] V. Dubée et al., ‘β-Lactamase inhibition by avibactam in Mycobacterium abscessus’, Journal of Antimicrobial Chemotherapy, vol. 70, no. 4, pp. 1051–1058, Sep. 2014, doi: 10.1093/jac/dku510.

[28] I. Halloum et al., ‘Deletion of a dehydratase important for intracellular growth and cording renders rough Mycobacterium abscessus avirulent’, Proc Natl Acad Sci U S A,vol. 113, no. 29, pp. E4228–E4237, Jul. 2016, doi: 10.1073/PNAS.1605477113/SUPPL_FILE/PNAS.201605477SI.PDF.

[29] J. C. van Kessel and G. F. Hatfull, ‘Efficient point mutagenesis in mycobacteria using single-stranded DNA recombineering: Characterization of antimycobacterial drug targets’, Mol Microbiol, vol. 67, no. 5, pp. 1094–1107, 2008, doi: 10.1111/j.1365-2958.2008.06109.x.

[30] E. Carbonnelle et al., ‘MALDI-TOF mass spectrometry tools for bacterial identification in clinical microbiology laboratory’, Clin Biochem, vol. 44, no. 1, pp. 104–109, 2011, doi: 10.1016/j.clinbiochem.2010.06.017.

[31] F. Ripoll et al., ‘Non mycobacterial virulence genes in the genome of the emerging pathogen Mycobacterium abscessus’, PLoS One, vol. 4, no. 6, Jun. 2009, doi: 10.1371/JOURNAL.PONE.0005660.

[32] K. A. Cohen, A. L. Manson, C. A. Desjardins, T. Abeel, and A. M. Earl, ‘Deciphering drug resistance in Mycobacterium tuberculosis using whole-genome sequencing: Progress, promise, and challenges’, Genome Med, vol. 11, no. 1, pp. 1–18, 2019, doi: 10.1186/s13073-019-0660-8.

[33] R. Mahajan, ‘Bedaquiline: First FDA-approved tuberculosis drug in 40 years’, Int J Appl Basic Med Res, vol. 3, no. 1, p. 1, 2013, doi: 10.4103/2229-516x.112228.

[34] N. J. Ryan and J. H. Lo, ‘Delamanid: first global approval’, Drugs, vol. 74, no. 9, pp. 1041–1045, 2014, doi: 10.1007/S40265-014-0241-5.

[35] G. Theron et al., ‘Evaluation of the Xpert MTB/RIF assay for the diagnosis of pulmonary tuberculosis in a high HIV prevalence setting’, Am J Respir Crit Care Med, vol. 184, no. 1, pp. 132–140, Jul. 2011, doi: 10.1164/rccm.201101-0056OC.

[36] N. K. Cuong, N. B. Ngoc, N. B. Hoa, V. Q. Dat, and N. V. Nhung, ‘GeneXpert on patients with human immunodeficiency virus and smear-negative pulmonary tuberculosis’, PLoS One, vol. 16, no. 7, Jul. 2021, doi: 10.1371/JOURNAL.PONE.0253961.

[37] M. G. B. F. de Faria et al., ‘Effectiveness of GeneXpert^®^ in the diagnosis of tuberculosis in people living with HIV/AIDS’, Rev Saude Publica, vol. 55, 2021, doi: 10.11606/S1518-8787.2021055003125.

[38] J. v. Philley et al., ‘Preliminary Results of Bedaquiline as Salvage Therapy for Patients With Nontuberculous Mycobacterial Lung Disease’, Chest, vol. 148, no. 2, pp. 499–506, Aug. 2015, doi: 10.1378/CHEST.14-2764.

[39] O. Lamrabet and M. Drancourt, ‘Genetic engineering of Mycobacterium tuberculosis: A review’, Tuberculosis, vol. 92, no. 5, pp. 365–376, 2012, doi: 10.1016/j.tube.2012.06.002.

[40] C. Chan, P. Pham, P. C. Dedon, and T. J. Begley, ‘Lifestyle modifications: Coordinating the tRNA epitranscriptome with codon bias to adapt translation during stress responses’, Genome Biol, vol. 19, no. 1, pp. 1–11, Dec. 2018, doi: 10.1186/S13059-018-1611-1/FIGURES/3.

[41] Y. H. Chionh et al., ‘TRNA-mediated codon-biased translation in mycobacterial hypoxic persistence’, Nat Commun, vol. 7, no. 1, pp. 1–12, Nov. 2016, doi: 10.1038/ncomms13302.

[42] World Health Organization, Global Tuberculosis Report 2020. 2020. Accessed: Nov. 13, 2020. [Online]. Available: http://apps.who.int/bookorders.

[43] V. C. N. de Moura, S. Gibbs, and M. Jackson, ‘Gene replacement in mycobacterium chelonae: Application to the construction of porin knock-out mutants’, PLoS One,vol. 9, no. 4, Apr. 2014, doi: 10.1371/journal.pone.0094951.

